# Aberrant resting-state functional brain networks in dyslexia: Symbolic mutual information analysis of neuromagnetic signals

**DOI:** 10.1101/272567

**Authors:** Stavros I. Dimitriadis, Panagiotis G. Simos, Jack M. Fletcher, Andrew C. Papanicolaou

## Abstract

Neuroimaging studies have identified a variety of structural and functional connectivity abnormalities in students experiencing reading difficulties. The present study adopted a novel approach to assess the dynamics of resting-state neuromagnetic recordings in the form of symbolic sequences (i.e., repeated patterns of neuromagnetic fluctuations within and/or between sensors).

Participants were 25 students experiencing severe reading difficulties (RD) and 27 age-matched non-impaired readers (NI) aged 7-14 years. Sensor-level data were first represented as symbolic sequences in eight conventional frequency bands. Next, dominant types of sensor-to-sensor interactions in the form of intra and cross-frequency coupling were computed and subjected to graph modeling to assess group differences in global network characteristics.

As a group RD students displayed predominantly within-frequency interactions between neighboring sensors which may reflect reduced overall global network efficiency and cost-efficiency of information transfer. In contrast, sensor networks among NI students featured a higher proportion of cross-frequency interactions. Brain-reading achievement associations highlighted the role of left hemisphere temporo-parietal functional networks, at rest, for reading acquisition and ability.

**Highlights:**

- Symbolic dynamics of MEG time series revealed aberrant Cross Frequency Coupling in RD students
- Global efficiency and strength of Cross Frequency Coupling could reliably identify RD students from age-matched controls
- Global Cost Efficiency, coupling strength, and the relative preponderance of cross-frequency interactions strongly correlated with reading achievement across groups.

## 1. Introduction

### 1.1. Cortical connectivity in reading and reading disability

There is accumulating evidence that the degree of myelination in left hemisphere cortico-cortical tracts correlates positively with reading skill (Hoeft et al., 2011; Niogi and McCandliss, 2006). Moreover, there is evidence (using Diffusion Tensor Imaging; DTI) of reduced myelination in left hemisphere white matter tracts connecting inferior frontal, temporal, occipital, and parietal regions among adults with a history of reading disability (Vandermosten et al., 2012). Both increased and decreased anatomical (using DTI) and functional connectivity (using task-related fMRI) within a network of dorsal and ventral brain regions have been reported in struggling readers compared to typically achieving readers (Richards et al., 2015). Other task-related fMRI studies reported reduced connectivity within the reading network in adults with a history of reading difficulties compared to non-impaired readers (Schurz et al., 2014; Van der Mark et al., 2011). fMRI evidence of a less integrated brain network has also been found in Chinese dyslexic children compared to typically achieving readers which was characterized by reduced long-range communication and increased local processing (Liu et al., 2015).

Studies on functional connectivity patterns at rest (i.e., independent of task performance) in dyslexia are scarce. Previous fMRI studies detected a strong association between functional connectivity in reading networks and reading ability in both children and adults (Koyama et al., 2011, 2013; Schurz et al., 2014; Zhang et al., 2014). Moreover, the strength of resting-state connectivity between the ventral visual word form area and the dorsal attention network was significant linked to individual reading skill (Vogel et al., 2014). Resting-state data may be particularly useful to assess aberrant modes of information exchange both within and between key reading-related cortical regions which may, in turn be associated with suboptimal cortico-cortical integration during reading acquisition and performance of reading tasks.

### 1.2. Cross-frequency coupling as a measure of resting-state functional connectivity

Magnetoencephalography (MEG) is uniquely suited to address functional connectivity because it possesses adequate temporal resolution to describe the real-time dynamics of fine-grained interactions between neuronal populations. By adopting a dynamic functional connectivity analysis of resting-state neuromagmetic data we identified abnormal temporal correlations between time series recorded over left temporo-parietal brain areas in students experiencing severe reading difficulties (RD) as compared to age-matched typical readers (Dimitriadis et al., 2013b). A more recent report using resting-state data from the same cohort focussed on cross-frequency coupling between neuromagnetic time series (Dimitriadis et al., 2016c). Sensor interactions in the form of phase-to-amplitude coupling were quantified though the phase-locking index which is thought to represent the degree to which slower brain rhythms in a given neuronal population can reset ongoing neurophysiological processes in a different neuronal population operating at higher frequencies (Buzsáki, 2010; Canolty and Knight, 2010; Buzsáki et al., 2013). Results indicated that resting-state activity in typical readers was characterized by more stable cross-frequency interactions than in RD students. One interpretation of these findings is that temporally stable cross-frequency information exchange reflects a typical and, presumably, optimal working level ensuring efficient neuronal transmission (Deco and Corbetta, 2011; Deco et al., 2013) available to support typical reading acquisition and performance.

In this study we extend these findings by examining both same-frequency and cross-frequency interactions in the same cohort of RD and typical readers. The novelty of the present report entails computing sensor interaction metrics based on the concept of *symbolic dynamics,* wherein neuromagnetic signals are first transformed into symbolic sequences consisting of a finite set of substrings (Janson et al., 2004; Dimitriadis et al., 2012a; Robinson and Mandell, 2016). Sensor interactions were then quantified using a variant of Mutual Information (King et al., 2013; Robinson and Mandell, 2016), a rather popular approach in the search for aberrant patterns of functional connectivity based on EEG and MEG recordings in a variety of clinical conditions (Colclough et al., 2017; Uhlhaas and Singer, 2006). The original Mutual Information algorithm was adapted here to accommodate symbolic time series and to compute an undirected weighted connectivity estimator (i.e., Symbolic Mutual Information). Surrogate data analyses were then used to identify the dominant type of intra-or cross-frequency coupling for each pair of sensors and construct a weighted, integrated functional connectivity graph characteristic of the resting-state recordings of each participant. Finally, estimated functional networks were spatially filtered through bootstrapping and submitted to graph analyses in order to assess both sensor-specific and overall network efficiency and cost-efficiency (Stam, 2014).

The present study had three aims: First, to identify aberrant spectral (intra-and cross-frequency coupling) and spatial characteristics of functional brain networks in RD students; Second, to assess the value of features associated with sensor-level brain network metrics in discriminating between RD and age-matched typical readers using machine learning techniques. Finally, to establish the functional significance of these metrics for basic reading skills through correlational analyses. We hypothesized that RD children would demonstrate reduced efficiency of information flow compared to non-impaired readers and sensor interactions that operate predominantly in same-frequency oscillations. Conversely, cross-frequency interactions would be more prominent in typical readers and their relative predominance will serve as a significant predictor of basic reading skill.

## 2. Material and Methods

### 2.1 Participants

Participants were two age-matched groups of students aged 7-14 years. The RD group included 25 children (12 boys) with reading difficulties (RD group) as indicated by scores below the 16th percentile level (standard score of 85) on the Basic Reading composite index (Word Attack and Letter–Word Identification subtest scores of the Woodcock–Johnson Tests of Achievement-III; Woodcock et al., 2001; WJ-III). These scores are consistent with the presence of dyslexia and is lower than in previous studies (Rezaie et al., 2011; Simos et al., 2011) of this cohort because we focused on studying severely impaired children. They were recruited from a larger Grade 6–8 intervention study (Vaughn et al., 2010) involving students at-risk for further academic failure because they failed to pass the school-administered Texas Assessment of Knowledge and Skills (TAKS).

Twenty-seven children (9 boys) who had never experienced difficulties in reading (NI group) served as a comparison group. These students had standard scores >90 on the Basic Reading Composite (corresponding to the 36th percentile) and were recruited from the same schools as the RD group in an attempt to control for educational history, ethnicity, and SES factors. All participants had Full Scale IQ scores >80 on the Wechsler Abbreviated Scale of Intelligence (Wechsler, 1999).

Detailed psychoeducational and demographic data are provided in Table 1. The two groups were comparable on age, ethnicity, handedness (there was one left handed student in each group), and Performance IQ (p > 0.1 in all cases). As expected the RD group demonstrated significantly lower scores than the NI group on measures of reading, Verbal IQ and spelling. Additionally, participants were selected for inclusion in either group only if they had T scores below 65 on the Attention Problems scale of the Child Behavior Checklist (Achenbach, 1991), as indicators of low risk for ADHD (Chen et al., 1994). Written informed assent and consent forms were signed by all participating children and their parents or guardians, respectively. The study had been approved by the Health Science Center-Houston and University of Houston Institutional Review Boards.

### 2.2. MEG Recordings

Three minutes of continuous brain activity was acquired with a whole-head neuromagnetometer array (4-D Neuroimaging MagnesWH3600), consisting of 248 first-order axial gradiometer sensors, housed in a magnetically shielded chamber. Participants were placed in a supine position and instructed to keep their eyes closed during the recording. Data were collected at a sampling rate of 1017.25 Hz and bandpass filtered in the 0.1–200 Hz range.

**Table 1.**
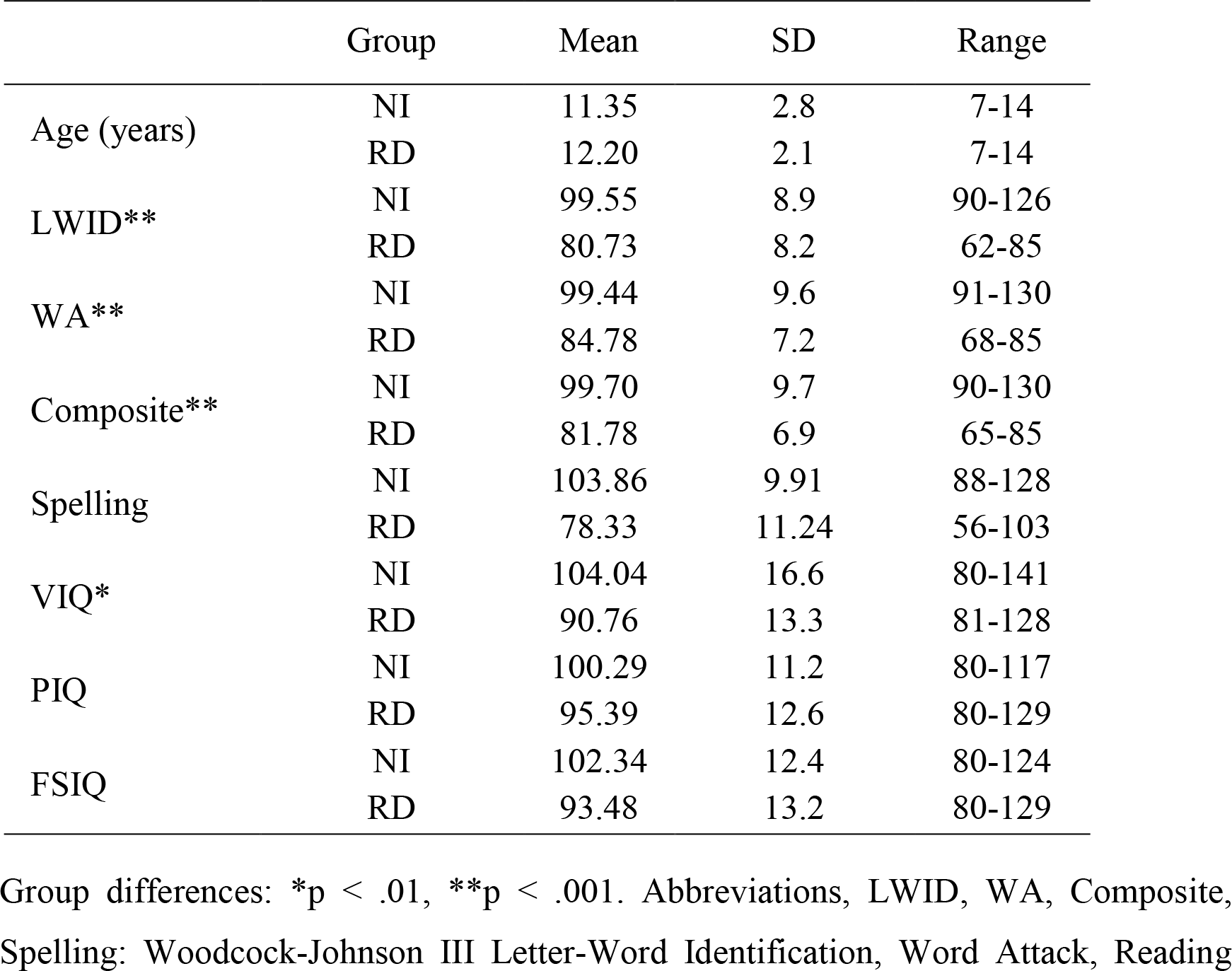
Demographic and psychoeducational data for typical (NI) and students with severe reading difficulties (RD).

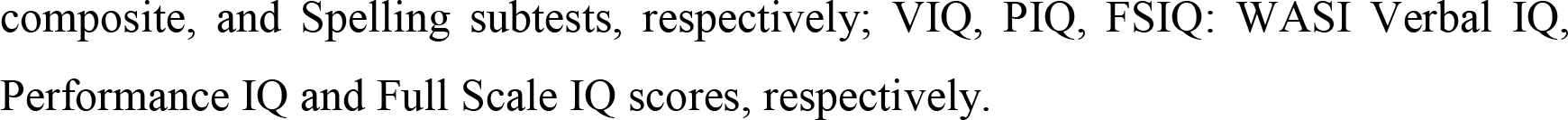

### 2.3 Data Preprocessing

The MEG data underwent artifact reduction using Matlab (The MathWorks, Inc., Natick, MA, USA) and Fieldtrip (Oostenveld et al., 2011). Independent component analysis (ICA) was used to separate cerebral from non-cerebral activity using the extended Infomax algorithm as implemented in EEGLAB (Delorme and Makeig, 2004). The data were whitened and reduced in dimensionality using principal component analysis with a threshold set to 95% of the total variance (Delorme and Makeig, 2004; Escudero et al., 2011). Kurtosis, Rényi entropy, and skewness values of each independent component were used to identify and remove ocular and cardiac artifacts. A given component was considered an artifact if, after normalization to zero mean and unit variance, more than 20% of its values were greater/lower than 2 SDs from the mean (Escudero et al., 2011; Dimitriadis et al., 2013a; Antonakakis et al., 2013, 2015). Additionally, we inspected the time course of each IC, its spectral profile, and the topological distribution of the IC weights to further confirm that an IC was an artifact. Across participants, the average number of ICs removed was 6 out of 50 ICs. The artifact-free ICs were then used to reconstruct the MEG time series. Axial gradiometer recordings were transformed into planar gradiometer field approximations using the sincos method implemented in Fieldtrip (Oostenveld et al., 2011). The data were then bandpass-filtered in the following frequency ranges using a 3^rd^-order Butterworth filter (in zero-phase mode): 0.5-4, 4-8, 8-10, 10-13, 13-15, 15-19, 20-29, and 30-45Hz corresponding to δ, θ, α1, α2, β1, β2, β3, and γ bands.

### 2.4 Spectral Power

For each participant and MEG sensor we calculated the power spectral density using the Fast Fourier Transform employing partially (50%) overlapping Hanning windows each consisting of 4096 data points. This yielded a frequency resolution of 0.25Hz. Relative power within each frequency band was calculated to assess the relative contribution of several oscillatory components to the global power (Leuchter et al., 1993; Rodriguez et al., 1999). The alpha peak frequency was also estimated to assess group differences. For further details on spectral power estimation see Supp. Material Sections 1-2.

### 2.5 Functional Connectivity indexed by Symbolic Mutual Information

Analyses described in this section aimed to assess relatively stable functional associations between every pair of MEG sensors. This procedure sought, first, to identify a finite set of systematic temporal patterns within each time series, reflecting the degree of signal predictability over time (the derived signal Complexity index is described in detail in Suppl. Material Section 4.1 and used as one of the comparison indices in participant classification as described in 2.10). Each pair of time series was then transformed into two symbolic sequences utilizing a *common* set of symbols (for more details see section 4.2 in Supp. Material). To achieve this goal the Natural Gas algorithm was adapted for pairs of time series (Dimitriadis et al., 2016a). The latter comprised of same-frequency/between-sensor pairs, cross-frequency/between-sensor pairs, and cross-frequency/within-sensor pairs. The strength of linear and non-linear functional associations for each pair of symbolic sequences was indexed by Symbolic Mutual Information (SMI), an undirected weighted connectivity estimator (King et al., 2013; Robinson and Mandell, 2015). SMI is defined as: 
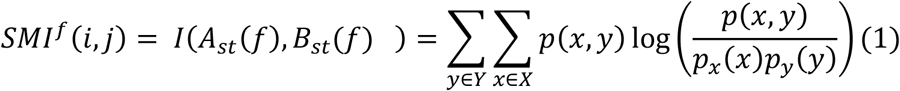
where 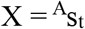 and 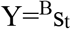, p(x,y)p(x,y) is the joint probability distribution function of X and Y and p_x_(x) = ∑_y∈Y_p(x, y) and p_y_(y) = ∑_x∈x_p(x, y) are the marginal probability distribution functions of X and Y, respectively. SMI values range between 0 and 1, with 0 denoting no functional coupling and 1 indicating perfect functional coupling over the entire recording period. This procedure resulted in a single *functional connectivity graph* per participant, frequency band (8), and pair of frequency bands (28) consisting of SMI values.

### 2.6 Dominant Intrinsic Coupling Modes for each pair of symbolic sequences

Individual functional connectivity graphs were further processed through surrogate data analyses to determine the Dominant Intrinsic Coupling Mode for each pair of symbolic sequences. 10,000 surrogate data sets were created by shuffling the symbolic sequence of the second MEG sequence 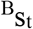 in each pair (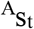 and 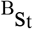) and re-estimated SMI values. Finally, a p-value was assigned to each pair of symbolic sequences (same-frequency/between-sensor, cross-frequency/between-sensor, and cross-frequency/within-sensor pairs) reflecting the proportion of surrogate SMI values that were higher than the observed SMI. This procedure produced a 3D tensor of p values for each participant of size 36 × 248 × 248. Significant Dominant Intrinsic Coupling Mode(s) for each pair of symbolic sequences were determined by applying a Bonferroni-adjusted statistical threshold of p < 0.01/36 = 0.00028 in order to control for family-wise Type I error. When more than one frequency or frequency pairs exceeded this threshold, the one associated with the lowest p value was retained. This procedure resulted in two 2D matrices for each participant of size 248 × 248: one containing the highest/statistically significant SMI values and the second the identity of the corresponding frequency or frequency pair (e.g., 1 for δ, 2 for θ, …, 8 for γ, 9 for δ-θ, …, 15 for δ-γ,…, 36 for β_3_-γ). In view of the purported importance of cross frequency interactions for efficient neuronal communication, the ratio of probability distributions of inter-frequency over the probability distribution of intra-frequency Dominant Intrinsic Coupling Modes was also computed (r index).

### 2.7 Topological Filtering and Graph Theory Analysis of Functional Brain Networks

In this stage of the analysis workflow, the integrated Functional Connectivity Graphs were submitted to topological filtering using surrogate data. This procedure relied on a data-driven variant of the Minimal Spanning Tree algorithm, namely Orthogonal Minimal Spanning Trees (OMST; for details of the algorithm see Dimitriadis et al., 2017a,b), and aimed to identify the subset of functional interactions that would ensure optimal information flow (indexed by network global efficiency) while minimizing the cost incurred by preserved functional connections. For a non-impaired reader, the GE-Cost vs. Cost function was optimized after 11 OMSTs leading to a selection of 2,689 out of 61,504 connections.

The relative importance of each sensor for information exchange within the individual functional connectivity graphs was quantified using the Global Efficiency metric derived from graph theory. Sensor-specific Global Efficiency was derived from the SMI values representing the Dominant Intrinsic Coupling Mode for each sensor pair and is defined as the inverse of the shortest distance between a given sensor and every other sensor. Network Global Efficiency reflects the overall efficiency of parallel information transfer within the entire set of 248 sensors and was estimated as the average sensor-specific GE value over all sensors using the following formula: 
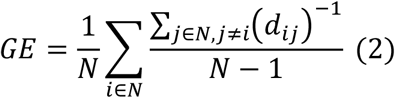

Overall Network Cost-Efficiency, defined as the global network efficiency minus overall network cost, was computed using a data-driven technique based on the maximization of overall network global efficiency: 
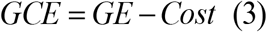

Cost was computed as the sum of the highest/significant functional connections divided by the sum of SMI values of the full-weighted functional network. Global Cost-efficiency was defined as the Global Efficiency at a given cost C minus the Cost value, which typically has a positive maximum value at some cost C_max_, for an economical small-world network. Importantly, this metric of network topology is independent of arbitrary, user-defined thresholds. Additionally, the Global Cost-Efficiency curve was estimated over a wide range of thresholds, and optimal threshold was derived for the maximum Global Efficiency value occurs at a specific Cost and Global Cost-Efficiency value (Bassett et al., 2009). Statistical group comparisons were conducted on overall network Global Efficiency, Cost, and Global Cost-Efficiency using the Wilcoxon Rank Sum Test.

### 2.9 Group-specific sensor subnetwoks

Dominant Intrinsic Coupling Modes that survived statistical (Section 2.6) and topological filtering (Section 2.7) were further processed using the Network Based Statistic toolbox (Zalesky et al., 2010) in an attempt to identify sets of sensor pairs forming a subnetwork, the strength of which significantly differed between the two groups. The nominal statistical threshold was set to 0.01, FDR-corrected over 5,000 iterations.

### 2.10 Classification of participants according to reading ability group based on network features

#### 2.10.1 Feature selection procedure

Initially, the optimal set of features to enter into the classification schemes were selected. Each original set of sensor-specific, univariate features (relative power, global efficiency, and signal complexity [see Supplementary Material Section 4.1]) consisted of 8 (frequency bands) x 248 (sensors) = 1984 features. Selection of optimal features from each set involved computation of Laplacian scores described in Section 1 of the Supplementary Material. Next, bootstrapping was employed to assign a p-value to each of the 1984 features. The criterion for feature selection was a Bonferroni-adjusted p < 0.05/(8 [frequency bands] * 248 [sensors]).

Derivation of features from the topologically-filtered Functional Connectivity Graphs involved dimension reduction using a procedure appropriate for multidimensional matrix data. For this purpose, we adopted Tensor Subspace Analysis (He and Cai, 2005), which treats the entire sensor network as a matrix (i.e., a network representation) rather than as a vector (i.e., a vectorized version of the weights comprising each graph). This method has been used successfully on resting-state (Dimitriadis et al., 2015a,b; Antonakakis et al., 2016) and task-related MEG data from mild traumatic brain injury patients (Dimitriadis et al., 2015b) in previous reports from our group. Tensor Subspace Analysis blends multi-linear algebra and manifold data learning. Given some Functional Connectivity Graph sampled from the space of functional connectivity patterns, the Tensor Subspace Analysis approximation is modeled by first building an adjacency graph capturing the proximity relationships among the connectivity patterns and then deriving a tensor subspace that faithfully represents these relationships. Tensor Subspace Analysis provides an optimal linear approximation to the Functional Connectivity Graph manifold. We selected a dimension of d=6 per dimension of the Functional Connectivity Graph which equals to a feature subspace of size 6 × 6.

#### 2.10.2 Classification procedure

Classification performance based on selected, sensor-specific relative power, symbolic complexity, and Global Efficiency features was assessed in separate schemes using two types of classifiers: a k–nearest neighbor algorithm and Support Vector Machines. Separate classification attempts were conducted on network-wise features, namely overall network Global Efficiency, Cost, Global Cost-Efficiency, and dimensions derived from the spatially-filtered weighted Integrated Connectivity Graphs through Tensor Subspace Analysis.

A 10-fold cross validation scheme was adopted each time. Each set of extracted features from the entire sample was randomly partitioned into two subsets, a *training set* (the database of features of known class) corresponding to 80% of the participants (20 NI and 22 RD students) and a *test set* (cases for which the class had to be predicted) corresponding to the remaining 20% of participants (5 NI and 5 RD students).

As a measure of classification performance we used the correct recognition rate which was calculated as the proportion of subjects in the test set for which the correct label was predicted. The cross-validation scheme was repeated 100 times and the mean value and standard deviation of the correct recognition rate, sensitivity, and specificity were estimated.

### 2.11. Associations between resting-state features and reading achievement

Potential predictors of Woodcock-Johnson III Basic Reading composite scores were assessed through correlational analyses performed on the entire sample of participants. The initial pool of features consisted of 248 sensor-specific Global Efficiency values, the SMI values of dominant coupling modes, where significant group differences were revealed through the Network Based Statistic, the overall network Global Efficiency, network cost, and network cost-efficiency values, and the r index. All indices had values ranging between 0 and 1. The measure of association used to construct the original correlation matrix between MEG-derived features and reading achievement scores was Distance Correlation which permitted quantification of linear as well as non-linear associations between every pair of features across groups (R; Szekely et al., 2007). The R metric ranges between 0 and 1 and has the important property that R(X,Y)=0 if and only if X and Y are truly independent. In order to control for multicolinearity among predictor variables, the original correlation matrix was reduced to a smaller number of feature clusters using a dominant-sets graph clustering algorithm (Dimitriadis et al., 2009, 2012a-d). In this method, the feature with the highest correlation coefficient with WJ-3 scores was retained from each of the remaining feature clusters.

Given the small number of participants compared to the number of independent variables, a leave-one out cross-validation scheme within an Extreme Learning Machine approach was followed to obtain multiple regression analysis results. Extreme Learning Machines have been shown to be suitable to handle difficult tasks without demanding extensive training sessions (Huang et al., 2006). They are feedforward Artificial Neural Networks with a single layer of hidden nodes, where the weights connecting inputs to hidden nodes are randomly assigned and never updated (Huang et al., 2015). In the current analyses, the Extreme Learning Machine was trained on MEG features and WJ-3 scores from N-1 participants to predict the scores of the Nth participant. This procedure was repeated N times. Fig. 1 summarizes the main steps of the proposed analytic procedure.

**Figure 1.**
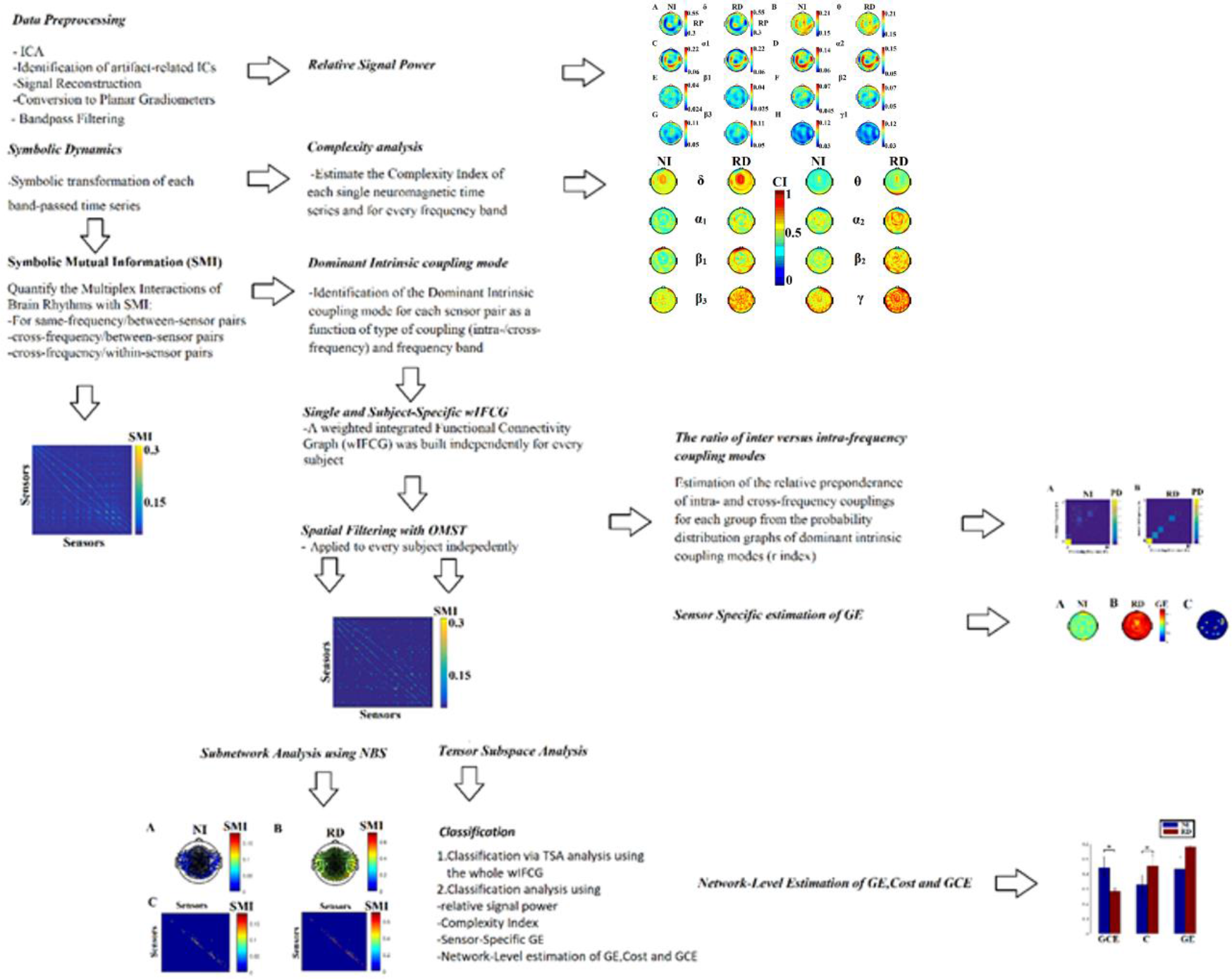
Data analysis flowchart. The analysis workflow and interdependencies among the different analysis procedures and computed metrics. Abbreviations, GE: Global Efficiency; GCE: Global Cost-Efficiency; ICA: Independent Components Analysis; NBS: Network-Based Statistic; OMST: Orthogonal Minimal Spanning Trees.

## 3 Results

### 3.1 Group differences on network efficiency and type of sensor-interactions

As a group RD students displayed lower sensor-specific global efficiency than NI students as shown in Figure 2a-b. These differences reached significance (p < 0.00001; Wilcoxon Rank Sum Test) for sensors located over left temporal, and bilateral parietal and frontal sensors (see Figure 2c). With respect to overall network metrics, RD students as a group displayed lower Global Efficiency and Global Cost Efficiency, and higher average Cost values as compared to the NI group (p < 0.00001; Wilcoxon Rank Sum Test; Fig. 3), suggesting less efficient network communication with higher cost in the former.

**Figure 2.**
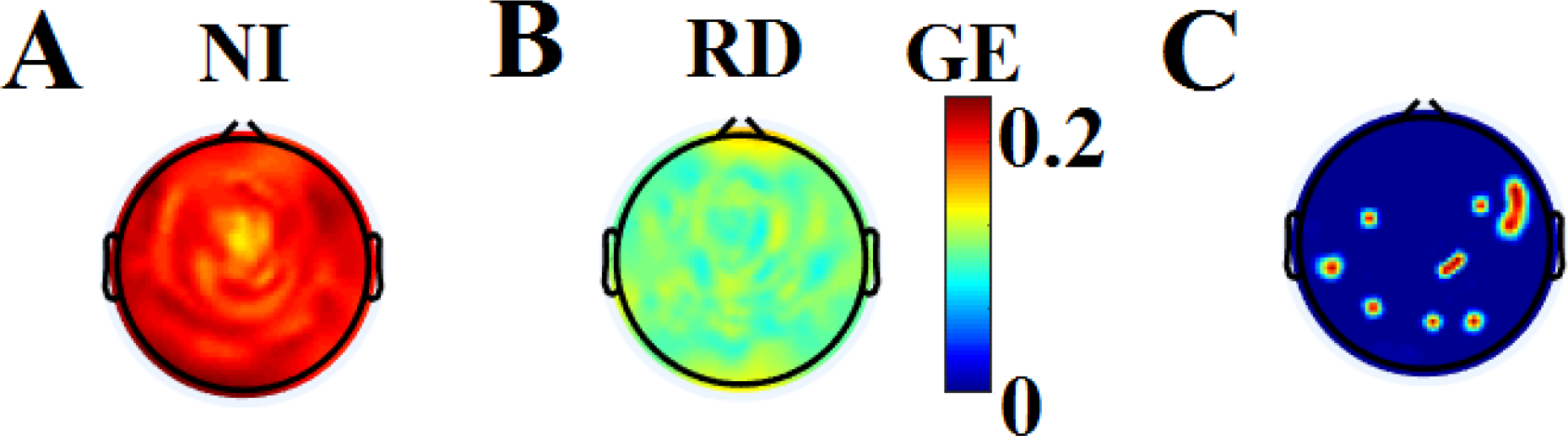
Group-averaged sensor-specific sensor-specific global efficiency (GE) for non-impaired (NI; A) and reading-disabled students (RD; B). Sensors where significant differences (NI>RD) were found are shown in red (p < 0.00001; Wilcoxon Rank Sum Test) (C).

**Figure 3.**
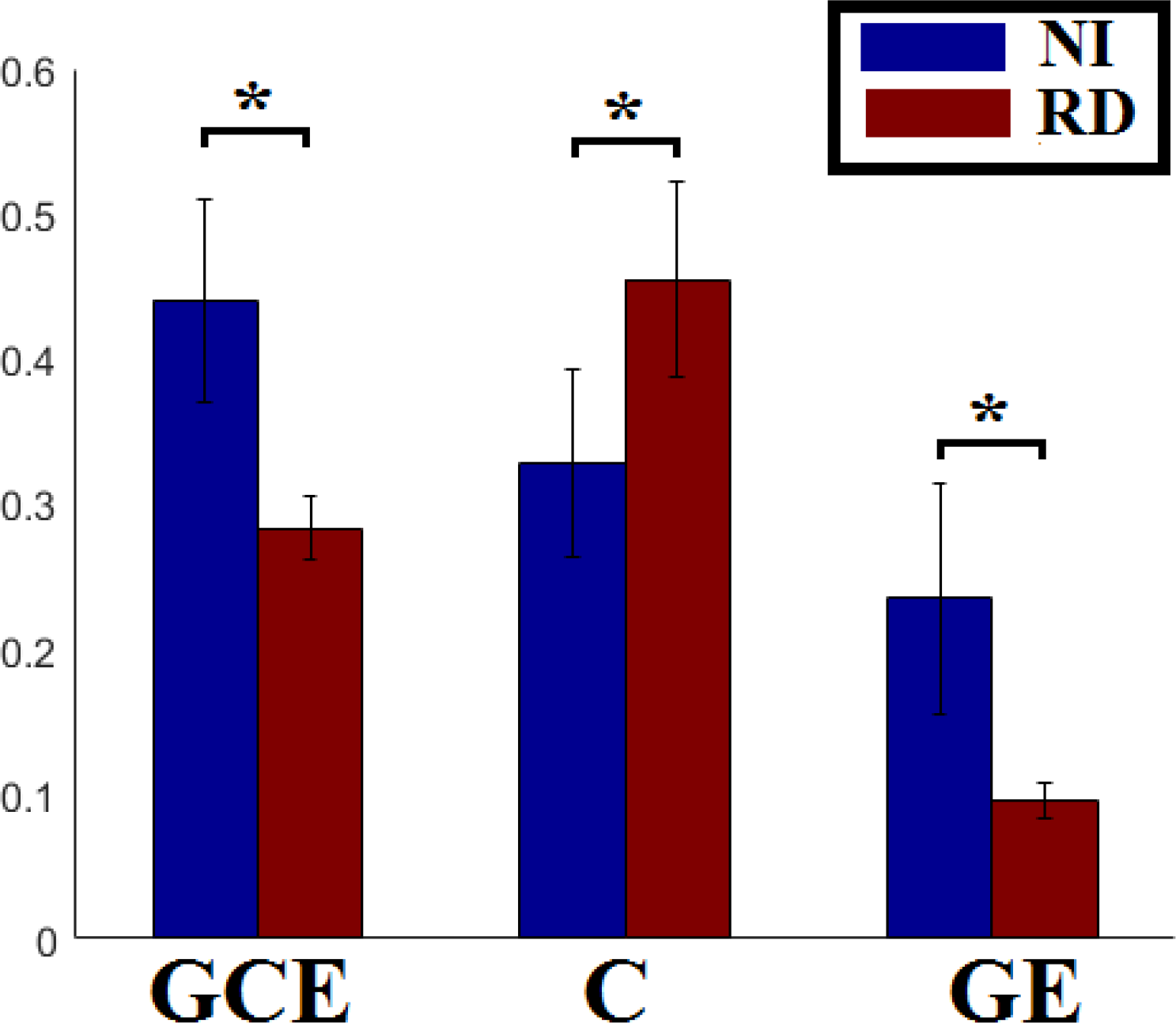
Group-averaged global cost-efficiency (GCE), cost (C) and overall network Global Efficiency (GE). Asterisks indicate significantly higher Cost values and lower GE and GCE values for Reading Disabled (RD) as compared to typical readers (NI; p < 0.00001; Wilcoxon Rank Sum Test).

The probability distributions of Dominant Intrinsic Coupling Modes for each group, computed across all sensor pairs, are shown in Figure 4. A notable finding is the higher percentage of significant cross-frequency coupling modes within among NI students (12%) compared to only 5% in the RD group. Conversely, RD students showed higher same-frequency probability distribution values in the θ, α1, and β1 bands compared to non-impaired readers. Interestingly, both groups showed prominent Dominant Intrinsic Coupling Modes in the δ band.

**Figure 4.**
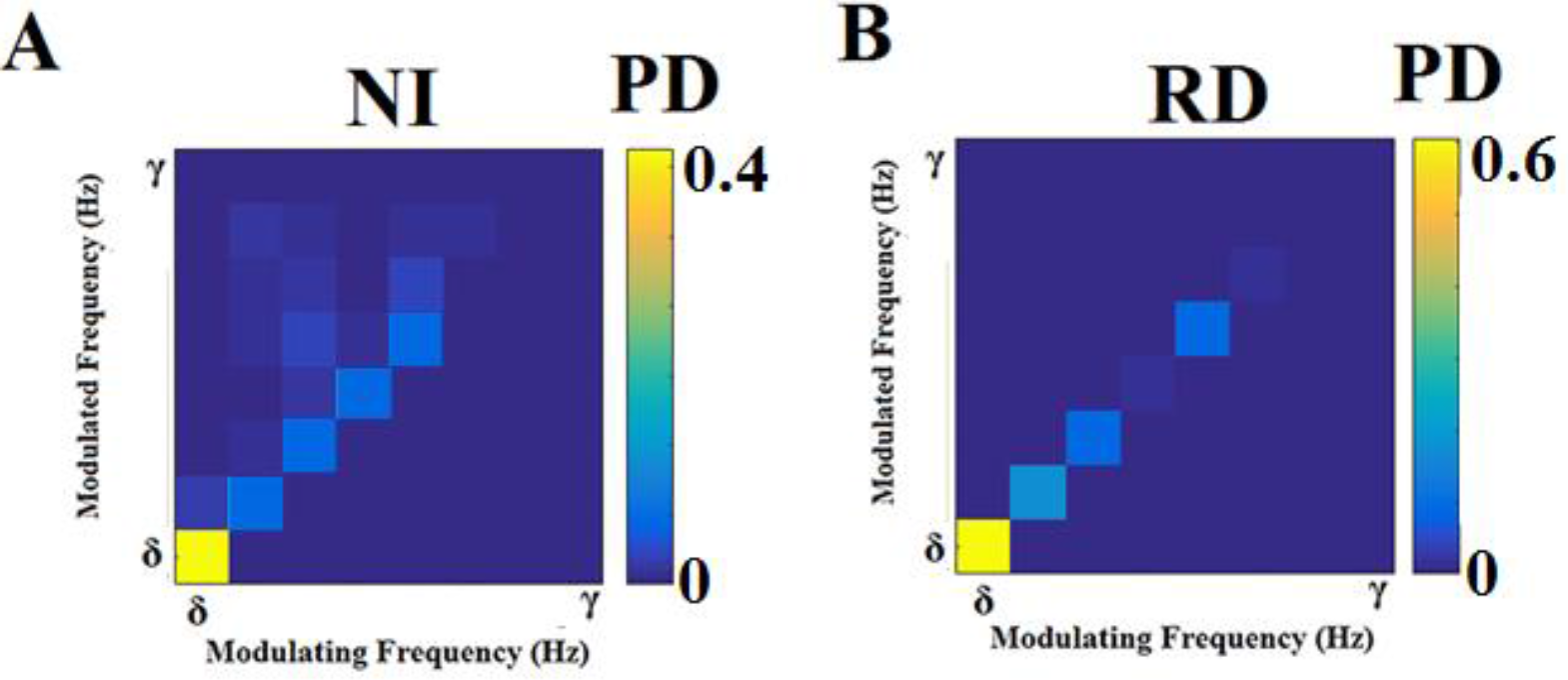
Group-averaged empirical Probability Distributions (PD) of dominant intrinsic coupling modes for NI (A) and RD (B) participants. The diagonal cells correspond to intra-frequency couplings; off-diagonal cells indicate cross-frequency interactions.

These results were corroborated by group comparisons at the subnetwork level. Specifically, significant group differences on SMI values based on the Network Based Statistic were detected for 537 sensor pairs which represented predominantly within-frequency interactions as illustrated in Fig. 5d. The strength of these interactions which are visualized in Fig. 5a-b was higher in the RD as compared to the NI group. Moreover, these interactions took place between neighboring sensors (given that nearly all significant SMI values were located at or near the diagonal in Fig. 5c). Analyses failed to reveal a subnetwork of sensor pairs featuring higher SMI values in NI as compared to RD students.

**Figure 5.**
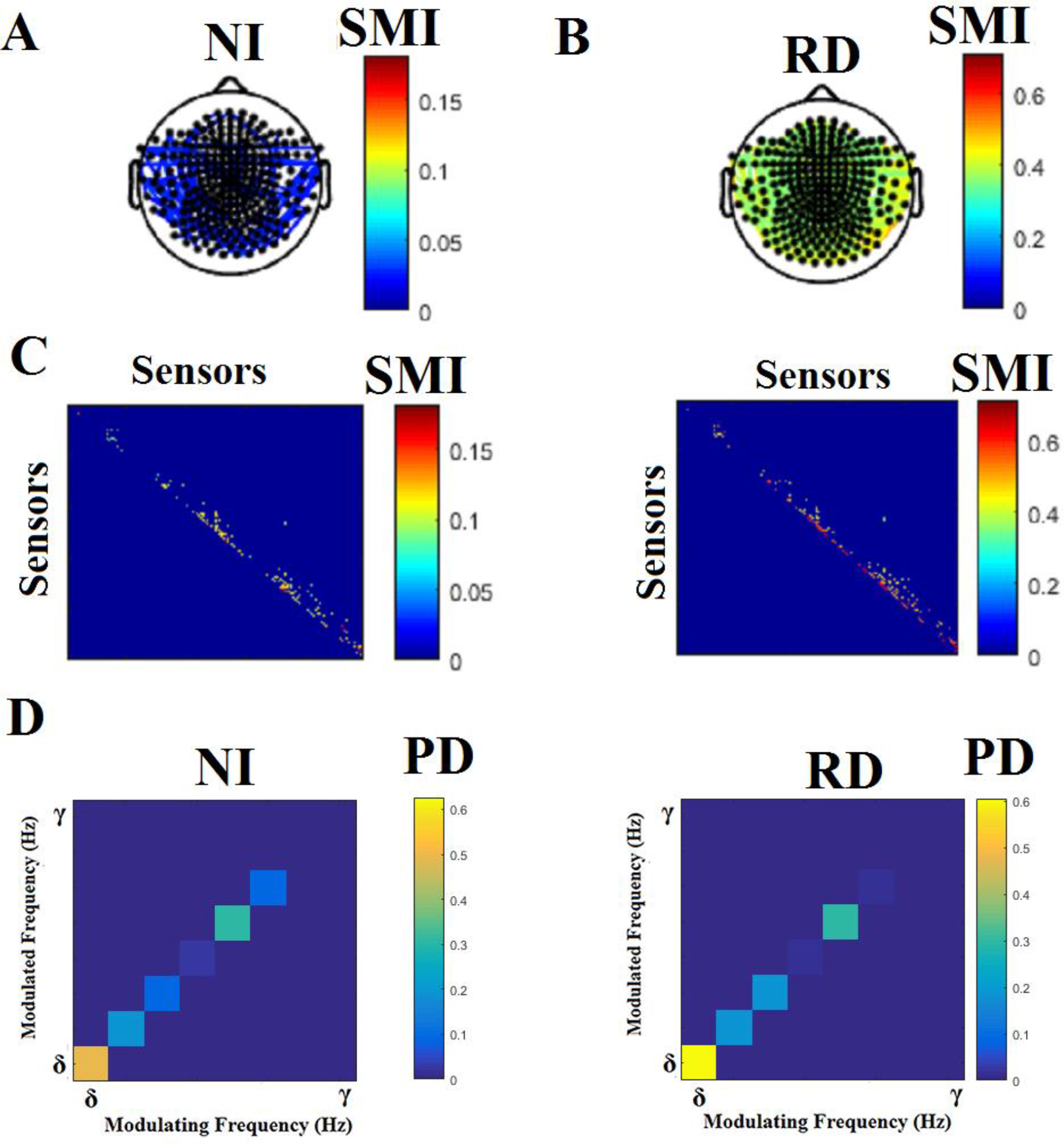
Group-characteristic dominant coupling modes based on Symbolic Mutual Information (SMI). The spatial layout of interactions that survived statistical (via bootstrapping) and topological filtering (via Orthogonal Minimal Spanning Trees) for each group of participants is illustrated in plots A-B. The vast majority of connections characterized by higher SMI values in the RD vs. NI groups were between neighboring sensors. This trend is visualized in the middle row of images (C) where SMI values are plotted as matrices with dimensions equal the number of MEG sensors. (D) Comodulograms shown in the lower row of images (D) indicate the dominant intra or cross-frequency couplings for each group.

### 3.2 Classification of students according to reading achievement groups

Table 2 reveals that overall classification accuracy using optimized sets of univariate features did not exceed 80% for symbolic Complexity and 70% for Relative Power (RP) measures (see Section 1 in Supplementary Material). Substantially higher classification rates were achieved using sensor-specific Global Efficiency (approximately 94%). Classification accuracy was slightly higher based on the 36 features derived from the topologically-filtered Functional Connectivity Graphs using Tensor Subspace Analysis. By comparison, classification accuracy based on overall network global Efficiency, Cost, or Global Cost-Efficiency did not exceed 60%.

**Table 2.**
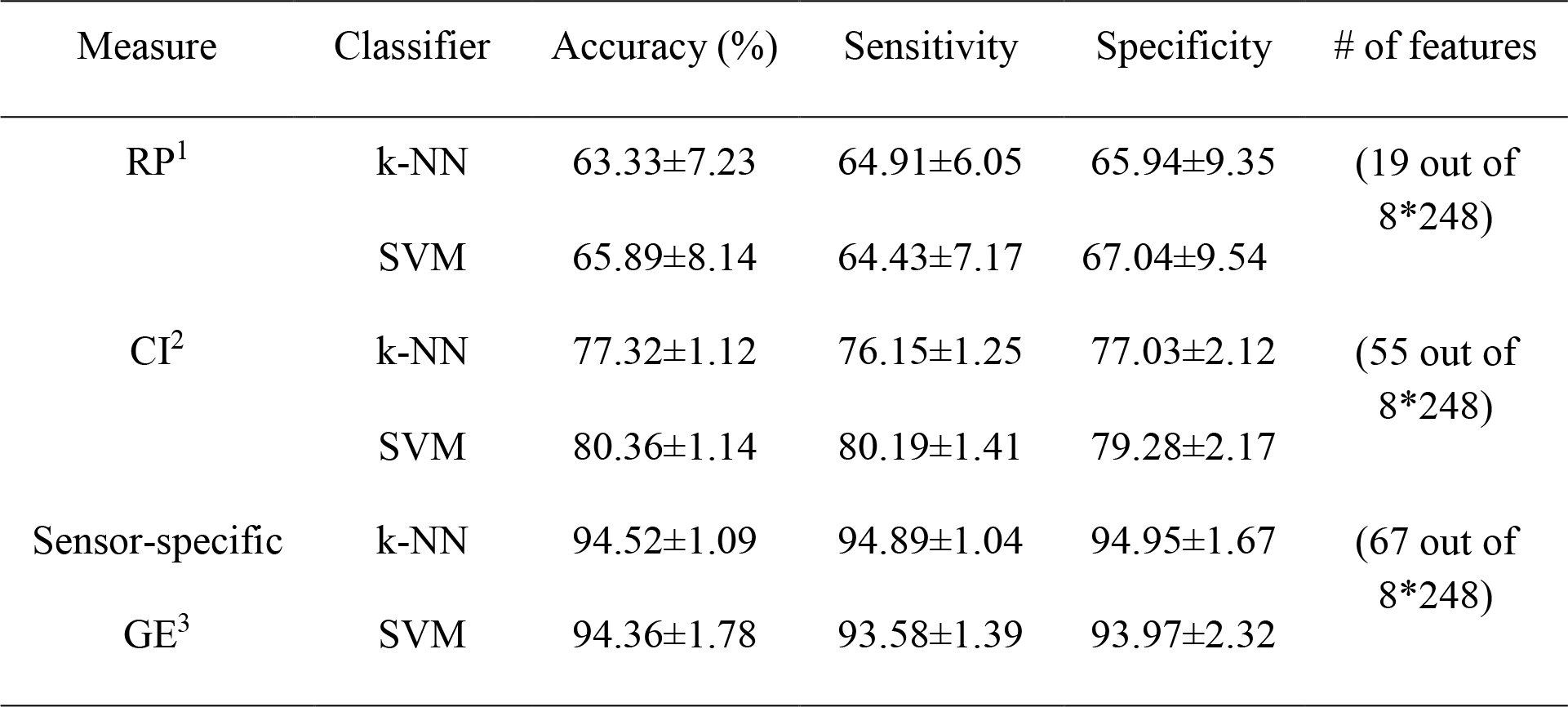
Results of various classification schemes using sensor-specific and network-level measures.

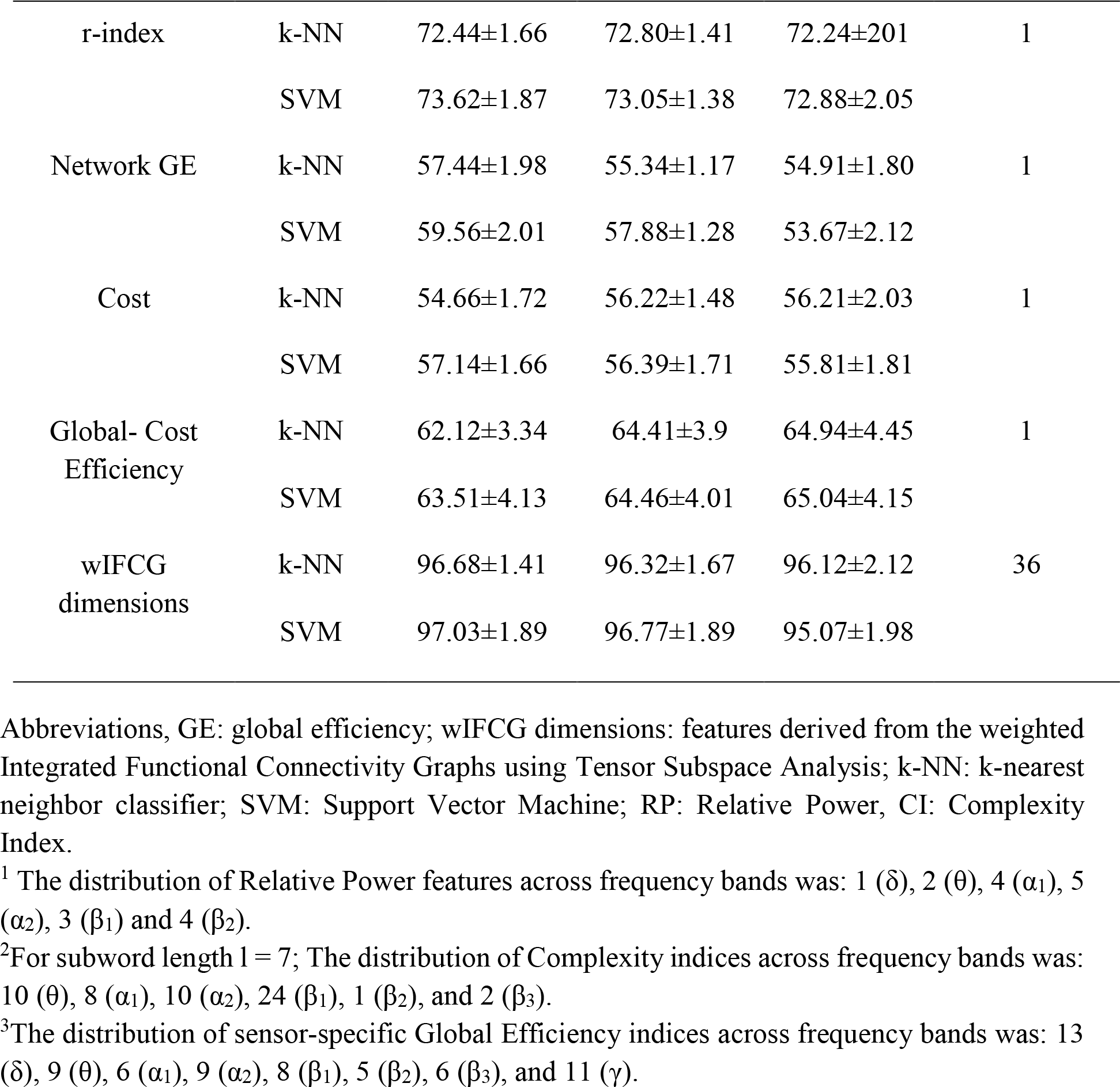

### 3.3 Predictors of reading achievement

The dominant-sets graph clustering algorithm reduced the original set or resting-state sensor-specific and network-related features to 63 feature clusters demonstrating the highest correlation coefficients with reading achievement scores. These features were input to an Extreme Learning Machine approach was implemented which was trained on data from N-1 participants to predict reading achievement scores of the Nth participant. The average R^2^ across N runs was 0.891. The set of features demonstrating the highest associations with reading achievement scores included 55 weighted sensor-pair interactions (SMI values), the r index, and Global Cost-Efficiency. Average correlation coefficients (±SD) between distinct types of features and WJ-3 scores were: r = 0.43±0.03 for SMI values, r = 0.47±0.03 for the r index, and r = 0.35±0.04 for network Global Cost-Efficiency. As a further cross-validation step of this approach we created 10,000 data sets by randomly selecting sets of 63 features (with replacement) and computed the compound R^2^ values obtaining an averaged R^2^ of 0.345 (SD = 0.067).

Figure 6 illustrates the spatial layout of the 53 sensor interactions which emerged as features with the highest association with reading achievement scores in the entire sample. In the RD group all weighted interactions were between temporo-parietal sensors in the α1 band (within-frequency), whereas corresponding interactions in the NI group were both within-frequency (in the α1 band involving parieto-occipital sensors) and cross-frequency (between α1 and β1 bands involving temporal sensors).

**Figure 6.**
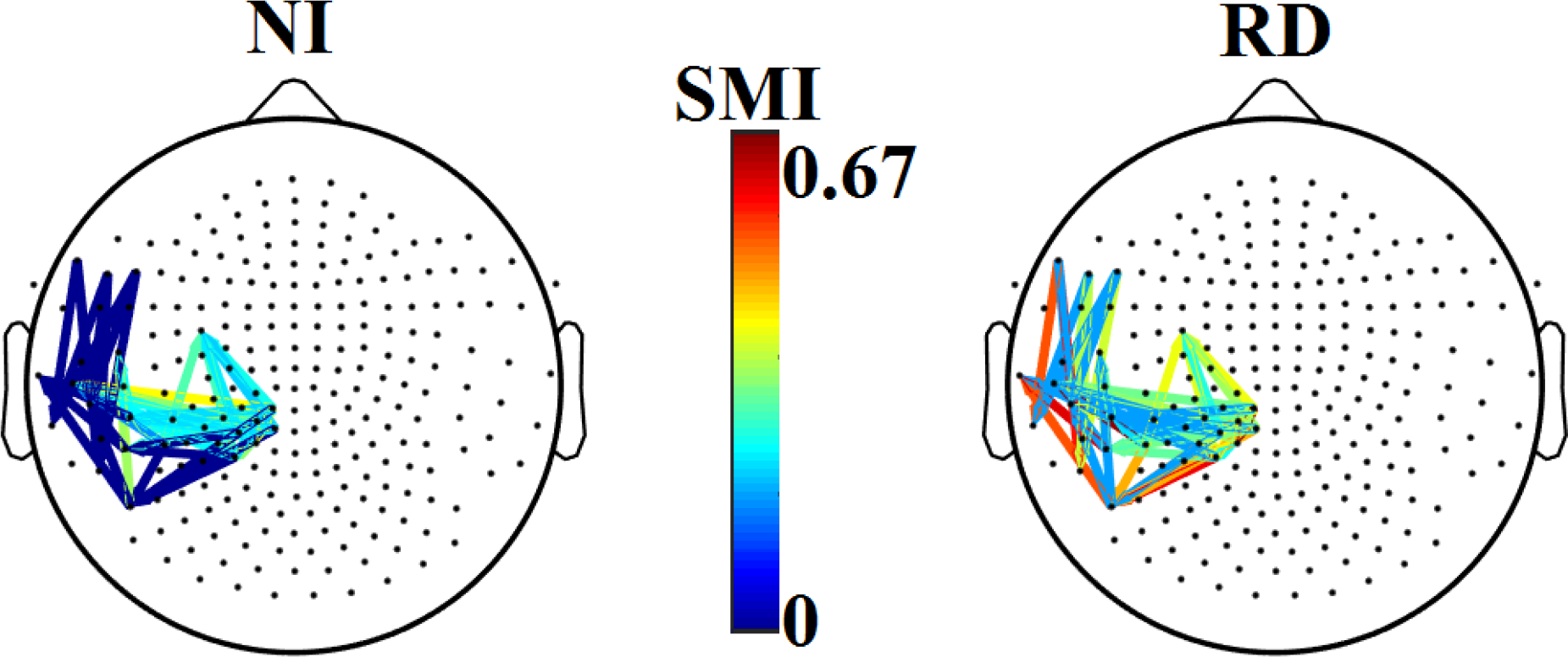
Topological layout of functional interactions that emerged as significant predictors of Woodcock-Johnson III Basic (WJ3) Reading composite scores for reading disabled (RD) and non-impaired readers (NI). The color scale indicates the strength (SMI value) of coupling for each of 53 pairs of sensors. Remarkably all interactions in the RD group were in the same frequency (a1 rhythm), whereas corresponding interactions in the NI group were both within-frequency (in the α1 band) and cross-frequency (between α1 and β1 bands).

## 4 Discussion

The present study explored a novel approach to represent systematic temporal variability of resting-state neuromagnetic time series as sequences of a finite set of distinct chunks of successive time points. Time series windows that displayed consistent time-varying profiles were represented by distinct symbols, with both symbol number and size determined empirically from the recorded data. A similar approach, adapted for sensor pairs, was utilized in order to detect common, repeatable patterns of symbolic sequences between sensors, providing a variety of indices of functional interaction. These included within and cross-frequency interaction for sensor pairs, sensor-specific global efficiency metrics derived from graph theory, as well as overall network efficiency and cost-efficiency in supporting information exchange over time.

Results can be summarized as follows: As a group, RD students displayed reduced global network efficiency and cost-efficiency of information transfer compared to non-impaired readers, which was predominantly realized through within-frequency interactions between neighboring sensors. In contrast, the repertoire of dominant sensor interactions among NI students featured a higher proportion (12% of total) of cross-frequency interactions, as compared to only 5% in the RD group.

The present findings extend previous results from our group revealing reduced local efficiency of short-range connections among left temporo-parietal sensors in the β3 band (Dimitriadis et al., 2013). Evidence suggesting a poorly integrated sensor-level network among children with dyslexia has also been reported based on minimal spanning tree analyses of resting-state EEG data (González et al., 2016). Although not directly comparable, these results are generally consistent with reports of disrupted network structure and various connectivity abnormalities in dyslexia (Frye et al., 2012; Finn et al., 2014; Koyama et al., 2010; 2013). Additional aberrant features of CFC interactions among RD students were highlighted in this study, representing globally reduced long-range CFC interactions compared to non-impaired readers. It has been proposed that cross-frequency interactions support the synchronization of nested hierarchical activities of neuronal assemblies oscillating on a dominant frequency mode (Buzsáki, 2010). This mechanism purportedly supports the accuracy in the timing of exchanged information among different oscillatory rhythms and the dynamic control of anatomically distributed functional networks (Buzsáki, 2006; Canolty and Knight, 2010).

The aforementioned group-level comparisons were supported by the results of individual classification schemes. Classification accuracy of individual students reached ~97% when network-level features were used, whereas classification accuracy relying solely on relative power at each sensor averaged 70%.

The significance of these results for the functional organization of the brain mechanism that supports basic reading skills was corroborated via correlational analyses. Results showed that an important positive predictor of Reading achievement scores was the relative preponderance of cross-frequency interactions both within and across-sensor pairs. Additional predictors were the strengths of symbolic interactions (both within and across-frequency) between several left temporo-parietal sensors in both groups of participants. These results are consistent with the widely held importance of left hemisphere networks for reading. Moreover, failure to identify short-range within-frequency interactions as predictors of reading achievement scores in the RD group suggest that the increased contribution of such interactions in the overall network of information exchange at rest represents a less efficient, compensatory mode of organization.

### 4.1 Methodological advances and limitations

The present results were facilitated by a number of methodological advances implemented in the current study. First and foremost, a novel approach was implemented in order to obtain a symbolic representation of continuous time series data. This symbolization scheme, which was first introduced in previous studies of our group (Dimitriadis et al., 2012; 2013), entails a data-driven procedure that determines the optimal symbol set size and the optimal symbol length to ensure representation of either individual time series or pairs of time series (Janson et al., 2004). Further, statistical and topological filtering were applied to the SMI-based functional connectivity graphs in order to identify the most prominent Dominant Intrinsic Coupling Mode features and to estimate both sensor-specific and overall network global efficiency and cost-efficiency. In the case of functional connectivity graphs this step was complemented by a tensorial approach which involves treating the data in its original matrix form preserving spatial associations between sensors. Additional features were also explored as sources of between-group variability in the form of relative power and global efficiency at each sensor, a step that had proven useful in increasing prediction accuracy of task-related EEG data in a previous study (Dimitriadis et al., 2015a).

The age-matched design of the present study did not permit us to assess whether the differences found between reading ability groups were, at least in part, due to group differences in rates of development of underlying brain mechanisms. The wide age range of the present sample may have introduced additional error variability in the estimation of reading achievement predictors (although standardized age-adjusted reading capacity scores were used in the analyses). Moreover, future studies should examine whether the pattern of functional network integration that is characteristic of RD children may change following systematic reading interventions. In the presence of change, do they represent compensatory processes or normalization towards a resting-state network that is similar to the one observed among typically achieving students? Another important limitation of the present findings concerns the nature of the MEG data employed in the analyses (sensor level), which significantly limited the spatial resolution of the results. It should also be noted that classification analyses were based on relatively small samples of RD and NI children and an extensive pool of MEG-derived features. Thus, despite efforts to select optimum independent variables to be used in the final classification models using bootstrapping, results may have still been largely inflated by the small subject to feature ratio. Independent validation of the final classification model in a different sample is necessary. Finally, it would be of considerable interest to explore how resting-state functional network organization is associated with similar features obtained during performance of reading tasks (e.g., Vourkas et al., 2011).

Finally, a cautionary statement is in order with respect to the generalizability of power spectrum data reported in the present work. Specifically, the strong similarity of the spatial distributions of average power across groups (as displayed in Fig S1-2) is notable. Both absolute and relative power are quantitative measures of brain activity integrated across experimental time. Conversion of axial gradiometer recordings into planar-gradiometer equivalent signals may have further contributed to this effect. More sophisticated approaches such as the Complexity Index introduced here and multiscale entropy that take into account the non-stationarity of brain activity may be more sensitive to group differences.

Additional precautions were taken in the present work to ensure that (a) group differences in the overall spectral profiles did not affect classification results (as indicated by comparable distributions of the n coefficient of the individual log[power[over log[frequency[functions shown in Fig. S3), and (b) the data submitted to the statistical and topological filtering and further used for group classification was not significantly contaminated by non-biological artifacts (using recordings empty room MEG recording shown in Fig. S5).

## 5 Conclusion

Reading-disabled children demonstrated a less efficient network communication compared to non-impaired readers characterized by reduced contribution of cross-frequency interactions between distant brain areas. The functional significance of these derived features was further supported by the linear prediction of Woodcock-Johnson III Basic Reading composite scores. The study relied heavily on the notion of Dominant Intrinsic Coupling Modes featuring both within and cross-frequency interactions, and on the optimal representation of signal dynamics as symbolic sequences to achieve very high rates of classification of individual students.

## Conflict of Interest Statement

The authors declare that the research was conducted in the absence of any commercial or financial relationships that could be construed as a potential conflict of interest.

## Acknowledgments

This research was supported in part by grant P50 HD052117 from the Eunice Kennedy Shriver National Institute of Child Health and Human Development (NICHD). The content is solely the responsibility of the authors and does not necessarily represent the official views of the NICHD or the National Institutes of Health. This research was also supported in part by a post-doc research fellowship to SD by the Greek State Scholarships Foundation (IKY).

